# Semi-supervised Encoding for Outlier Detection in Clinical Observation Data

**DOI:** 10.1101/334771

**Authors:** Hossein Estiri, Shawn N Murphy

**Affiliations:** Harvard Medical School, Boston, MA, USA; Massachusetts General Hospital, Boston, MA, USA

**Keywords:** Neural Networks, Encoding, Semi-supervised Encoding, Outlier Detection, Data Quality, Electronic Health Records

## Abstract

**Background and Objective:** To evaluate the utility of encoding for outlier detection in clinical observation data from Electronic Health Records (EHR).

**Methods:** This article presents a semi-supervise encoding approach (super-encoding) for constructing a non-linear exemplar data distribution from EHR data and detecting non-conforming observations as outliers. Two hypotheses are tested using experimental design and non-parametric hypothesis testing procedures to evaluate the outlier detection performance of the semi-supervised encoding approach and increasing demographic precision in encoding.

**Results:** The experiments involved applying 492 encoders to 30 laboratory tests extracted from the Research Patient Data Registry (RPDR) from Partners HealthCare. We report results obtained from 14,760 encoders. The semi-supervised encoders (super-encoders) outperformed conventional autoencoders in outlier detection. Adding age at observation to the baseline encoder (that only included observation value as the feature) slightly improved outlier detection. Top-nine performing encoders are introduced. The best outlier detection performance was from a semi-supervised encoder, with observation value as the single feature and a single hidden layer, built on one percent of the data and one percent reconstruction error. At least one encoder had a Youden’s J index higher than 0.9999 for all 30 observations.

**Conclusion:** Given the multiplicity of distributions for a single observation in EHR data (i.e., same observation represented with different names or units), as well as non-linearity of human observations, encoding offers huge promises for outlier detection in large-scale data repositories.

## Introduction

Data from Electronic Health Records (EHR) can accelerate real-time translation of evidence-based discoveries in everyday healthcare practice as we strive towards a rapid-learning health care systems.[1,2] However, truthfulness (plausibility) of observations stored in EHRs is can be disputable,[3] as these systems were not initially designed for research and discovery. For example, it is plausible to come across observation values in EHRs that are biologically implausible – e.g., a BMI of 1,400). It is extremely unlikely for such observation records to represent the “truth” about a patient at a given clinical encounter and at a given time. Systematic detection of such plausibility issues in EHR data are difficult for three reasons. First, although there are often gold-standards for high/low “normal” values, global gold-standards are not always available to set cut-off ranges for high/low “implausible” values. Second, even if such global cut-off ranges are available for all observations, they may not fully capture biologically implausible values due to an inherent non-linearity in observation data, as these values are expected to vary from patient to patient, for instance, by age, race, gender – i.e., a weight record of 700 lbs. may be plausible for a 50-year-old Male, but certainly not for a 5-month-old. Third, the multiplicity of observations and their representations in EHRs (i.e., same observation represented with different names or units), along with institutional variations in data standards are significant hurdles to systematic implementation of manual procedures to detect plausibility issues. Nevertheless, hard-coding cut-off ranges are the standard approach in many clinical data repositories for identifying implausible values.

With the abundance of unlabeled data (e.g., vital signs) in EHR repositories, unsupervised learning (a.k.a., predictive learning) offers promising applications in characterizing clinical observations into meaningful sub-groups that can embody non-linear properties. In particular, Neural Networks provide low-cost opportunities for constructing adaptive data representations that can be used to evaluate EHR data plausibility. We demonstrate the utility of feature learning by auto-encoders, as well as a semi-supervised encoding approach, for outlier detection in clinical observation data, including records of vital signs and laboratory result values.

## Background

There is no universally agreed upon definition for an outlier.[4] Generally, outlier detection refers to the problem of discovering data points in a dataset that do not conform to an expected exemplar.[5] Across different domains, such ‘non-conforming’ observations are referred to as, noise, anomaly, outlier, discordant observation, deviation, novelty, exception, aberration, peculiarity, or contaminant.[4,5] Outlier detection methods are often based on silver- or gold-standard ‘inlier’ data distributions, or statistical or geometric measures (e.g., Euclidean distance).[6]

Our approach aims to model an exemplar (i.e., inlier) distribution from the data that best represents the underlying patterns in the data, and utilize that distribution to benchmark any other observation, as inliers or outliers. In the context of clinical data, biologically implausible values are examples of observations that do not conform to the exemplar distribution of clinical observations, when massive loads of observation data are available. Our approach takes the view of letting the high-throughput EHR data speak for itself without relying on too many assumptions about what might be plausible or implausible.

We use Neural Networks for this purpose. The primary application of Neural Networks (NNs) is in supervised learning. If provided with the right type of data, deep Neural Networks are powerful learning algorithms with widespread applications across many domains.[7] However, application of supervised learning in general is dependent on presence of labeled input data, which are often manually generated.[8] This dependency limits application of deep Neural Networks in fields where much of the raw data are either still unlabeled or labeled data may not be reliable.

## Outlier Detection with Autoencoders

The central idea in Neural Networks is to model outputs as a non-linear function of linearly derived input features,[7] which will enable us to define complex forms of hypothesis *h_wb_*, given an activation function *h*, and parameters *w* (or weight) and *b* (or bias) between units in different layers.[8] NNs have been widely applied to anomaly detection tasks.[4,5,9]

Autoencoders (a.k.a., auto-associative Neural Networks) are unsupervised NN algorithms that automatically learn features from unlabeled data,[8,10] and are efficient for learning both linear and non-linear underlying characteristics of the data without any assumptions or a priori knowledge about input data distribution.[9] Autoencoders can learn non-linear patterns without complex computation.[11] Since their introduction in the 1980s, autoencoders have been successfully applied in the deep architecture approach, dimensionality reduction, image denoising, and information retrieval tasks.[10,12,13]

Suppose we have a vector of unlabeled data, {*x*^(1)^, *x*^(2)^), …}, where *x*^(*i*)^*∈* ℝ^(*n*)^. An autoencoder applies backpropagation to reconstruct the input values, by setting the target values (*y*^(*i*)^) to be equal to the input values (*x*^(*i*)^). In other words, the autoencoder aims to learn function *h_wb_*(*x*) that approximates 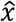 with the least possible amount of distortion from *x*. It compresses the input data into a low dimension latent subspace and decompresses the input data to minimize the reconstruction error, *ε*:[11]

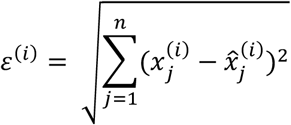

Autoencoders often have a multi-layer perceptron feed-forward network architecture in a bottleneck (Figure 1), consisting of an encoder and a decoder that produces 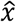 as a reconstruction of *x*.[10]

**Figure 1.**
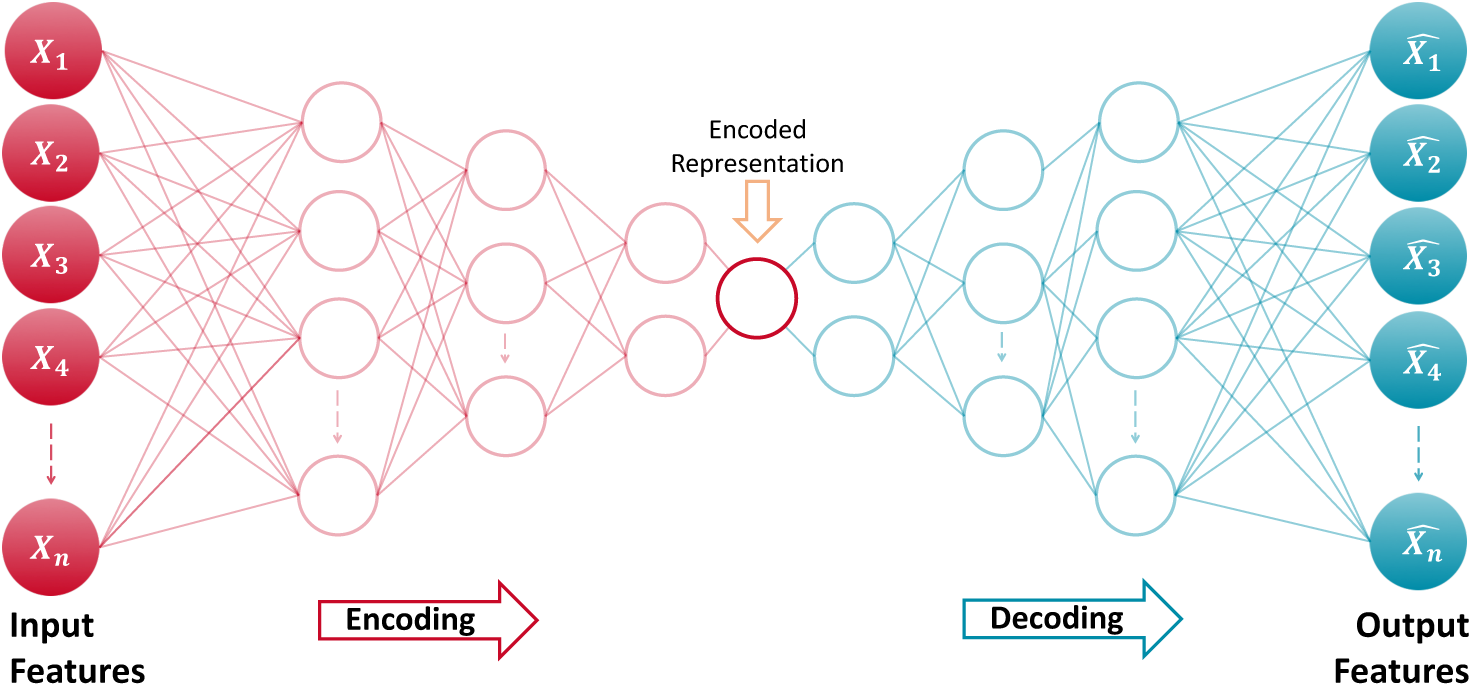
Architecture of an autoencoder

An autoencoder that perfectly reconstructs the input data (i.e., *h_wb_*(*x*) = *x*)) is not useful; instead, the goal is that function *h* takes on useful properties from the input data.[10] With this procedure, a worthwhile application of autoencoders is in outlier detection by identifying data points that have high reconstruction errors.[11] For example, Hawkins et al. (2002) used a 3-layer perceptron Neural Networks – also known as Replicator Neural Network (RNNs) – to form a compressed model of data for detecting outliers. [14] In anomaly detection with autoencoders, the reconstruction error is used as the anomaly score, where low reconstruction error values denote normal instances (or inliers) and large reconstruction errors represent anomalous data points (or outliers).[11]

Properly regularized autoencoders outperform Principal Component Analysis (PCA) or K-Means methods in learning practical representations and characterizing data distribution,[6] and are more efficient in detecting “subtle” anomalies than linear PCAs and in computation cost than kernel PCAs.[11,13]

## Hypotheses

We conduct experiments in which we evaluate the utility of encoding for the purpose of detecting outliers in clinical observation data from EHRs. We test two hypotheses. First, we hypothesize that increasing precision in training data by introducing demographic features stored in EHRs – using the i2b2[15,16] patient dimension – may lead to an even more precise learning of the exemplar function *h_wb_*(*x*) and improve outlier detection.

Given the large amount of clinical observation data in EHR repositories and the sheer ability of Neural Networks in learning data representations, we suspect that a conventional implementation of autoencoders – that uses the entire data for compression (i.e., encoding) – may result in a poor outlier detection performance, due to inclusion of noisy data in the learning process. Therefore, our second hypothesis is that training the encoder function on a small random sample of data (as a silver-standard distribution) may result in construction of a more precise “exemplar” representation of the underlying data distribution, and result in a more efficient outlier detection. We call this implementation semi-supervised encoding, or super-encoding.

## Data

To test these hypotheses, we used data on over 60 clinical observations, including records on laboratory results from Research Patient Data Registry (RPDR) from Partners HealthCare,[17] for which we were able to obtain global gold-/silver-standard cut-off ranges through literature search, and therefor calculate sensitivity and specificity for comparing the encoders (see Appendix A). For some of the laboratory tests, there was no record outside the implausible range, which would not allow us to calculate sensitivity scores. Removing those results, we report outcomes from 30 lab tests. The average dataset for each observation contained more than seven million observation records.

For each health record, we calculated patient-specific age at observation. We also created a joint demographic variable from Race and Ethnicity – a list and definition of demographic variables is available in Appendix B.

## Design

Although autoencoders are often trained using a single-layer encoder and a single-layer decoder, using deep encoders and decoders offer many advantages, such as exponential reduction in computational cost and amount of training data needed, and better compression.[10] Therefore, we built encoders with different NN architectures to allow for the an improved outlier detection performance.

To test our hypotheses, we developed 492 forms of hypotheses *h_wb_* per each of the 32 observations from combinations of seven sets of input features, 12 Neural Network architectures (i.e., patterns of connectivity between neurons), four sampling strategies (1%, 5%, 10% for super-encoding, and 100% for autoencoding), and three reconstruction error cut-off points for outlier detection. The following six steps describe the design of our experiments:

1. To test out first hypothesis, a vector of input features is selected from {*x*^(1)^, *x*^(2)^, … *x*^(12)^}, where, *x*^(1)^: observation value (OV) – the baseline feature; *x*^(2))^: age at observation (A); *x*^(3)^, *x*^(4)^: gender (G) – male or female; and *x*^(5)^ - *x*^(12)^: Race/Ethnicity (R/E).
2. A symmetrical bottleneck Neural Network architecture is selected to extract parameters in *h_wb_*. We restricted the choice of NN architecture such that the maximum depth of the network would not exceed the number of input features.
3. To test our second hypothesis, a sampling strategy is applied. A random sample of size *n* is drawn from the observation data with size *N*, where,

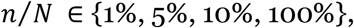

resulting in a vector of input features {*x*′^(1)^, *x*′^(2)^, … *x*′^(12)^}, based on the choice made in step 1 – *when n* = 100%, *then x*′ = *x* samples between 1-10% construct semi-supervised encoders, and 100% sample construct conventional design of autoencoders.
4. *h_wb_* is trained on *x*′, *h_wb_*(*x*′), with the goal of minimizing the reconstruction error for 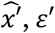:

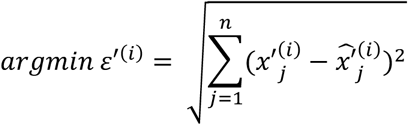
5. *h_wb_* is used on *x* to reconstruct 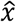 and calculate *ε*.
6. Outliers are flagged when *ε* ≥ *θ*, the error margin ratio, where,

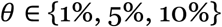

through the last step we allow *θ* vary by feature, NN architecture, and sampling strategy.

## Implementation

We used the autoencoder implementation in H2O [18] and developed functions to run the experiments – functions to run the experiments, results, and analytic code to process the results are available on GitHub (*link will be posted after the peer review process*). We also used *tanh* as the activation function. H2O normalizes the data to the compact interval of (−0.5, 0.5) for autoencoding to allow bounded activation functions like *tanh* to better reconstruct the data.[18]

## Performance Indices

Selecting an appropriate cut-off point to evaluate performance of AI algorithms depends on the context.[19] For each of the 492 encoders, we calculated the true-positive rate (TPR), a.k.a. sensitivity (Se) or recall, which represents the probability of truly detecting (Power) outliers, and false-positive rate (FPR) or probability of false alarm (Type I error) – FPR = 1 – specificity (Sp). Minimizing false positive rates while maximizing detection of true outliers are the main evaluation criteria for outlier detection methods.[9] To pick the best overall encoders, we calculated Youden’s J index[20] based on the following formula:

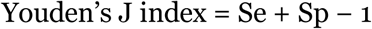

The goal was to maximize the Youden’d J index, and thereby maximize the combined specificity and sensitivity.

## Non-parametric hypothesis testing

To evaluate our hypotheses, we used non-parametric and post-hoc tests for comparing and ranking the outlier detection performances a cross the 14,760 encoders (492 encoders × 30 observations). These data points are more than enough for non-parametric hypothesis testing. The goal was to evaluate whether the Youden’d J indices would provide enough statistical evidence that the encoders have different outlier detection performances. Specifically, we applied Friedman post-hoc test,[21] with Bergmann and Hommel’s correction,[22] for comparing and ranking all algorithms.[23,24] Before performing the Friedman test, we ran Nemenyi’s test to perform an initial ranking, identify the Critical Difference (CD), and create a ranking diagram. The CD diagrams provide effective summarizations of encoder ranking, magnitude of difference between them, and the significance of observed differences. Any two algorithms who have an average performance ranking difference greater that the CD are significantly different.[24]

## Results

We compiled results of the 492 encoders on the 30 observations. Figure 2 illustrates the receiver operating characteristic (ROC) plots and Youden’d J indices for each observation – grouped by Logical Observation Identifiers Names and Codes (LOINC). LOINC provides common identifiers, names, and codes for health measurements, observations, and documents. The list of labs, their associated LOINC, and silver-standards cutoff ranges are provided in Appendix A.

**Figure 2.**
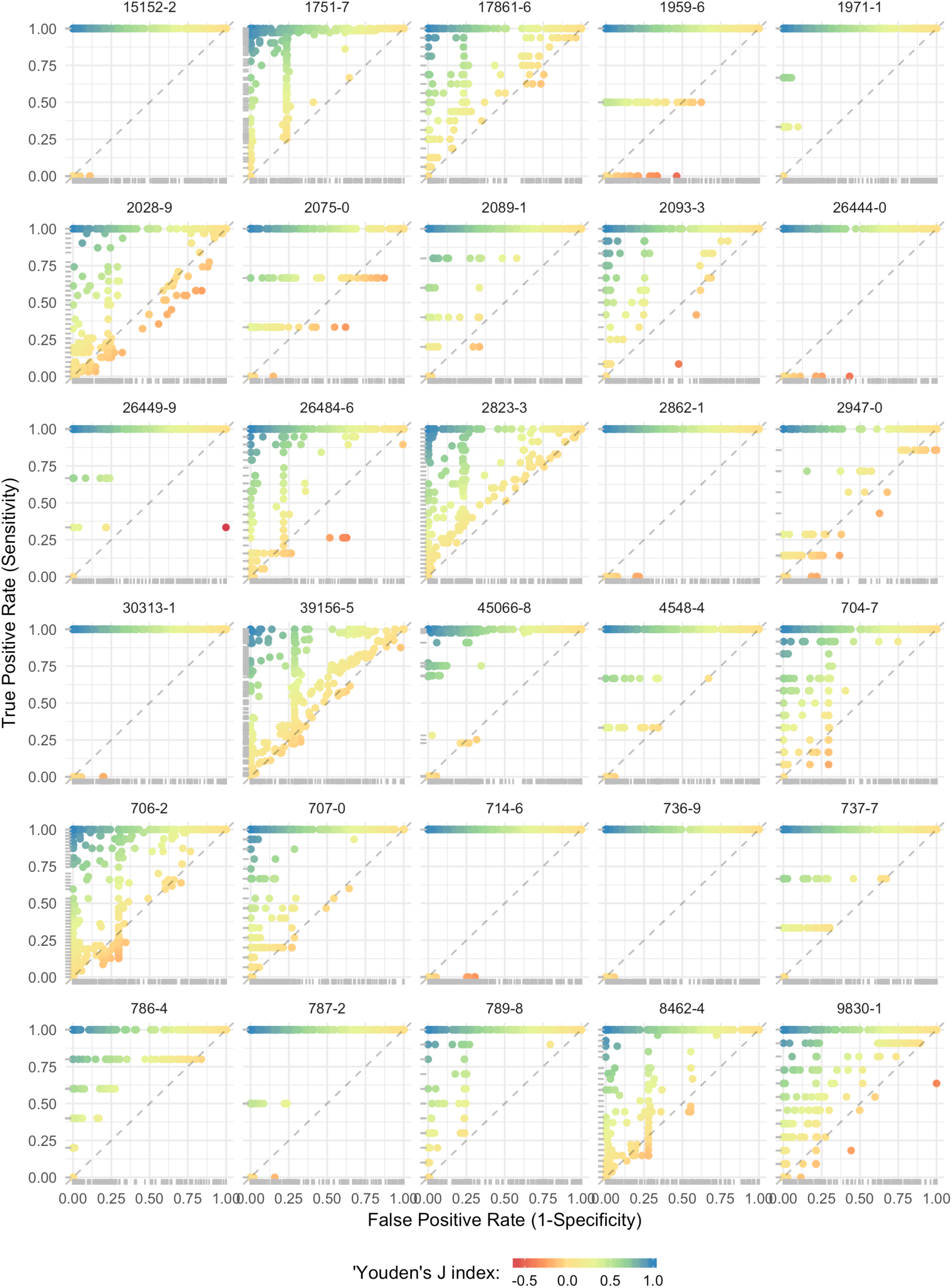
ROC plots and Youden’d J indices for each observation. * dots represent results from discrete encoders, so the ROC may not be interpreted as a curve.

### Testing the first hypothesis

To evaluate the first hypothesis, we performed non-parametric hypothesis testing on the entire encoders based on the features that formed them. Figure 3 presents the Critical Difference (CD) diagram for encoder features. On the CD diagram, each encoder is placed on the X axis according to its average performance ranking.

**Figure 3.**
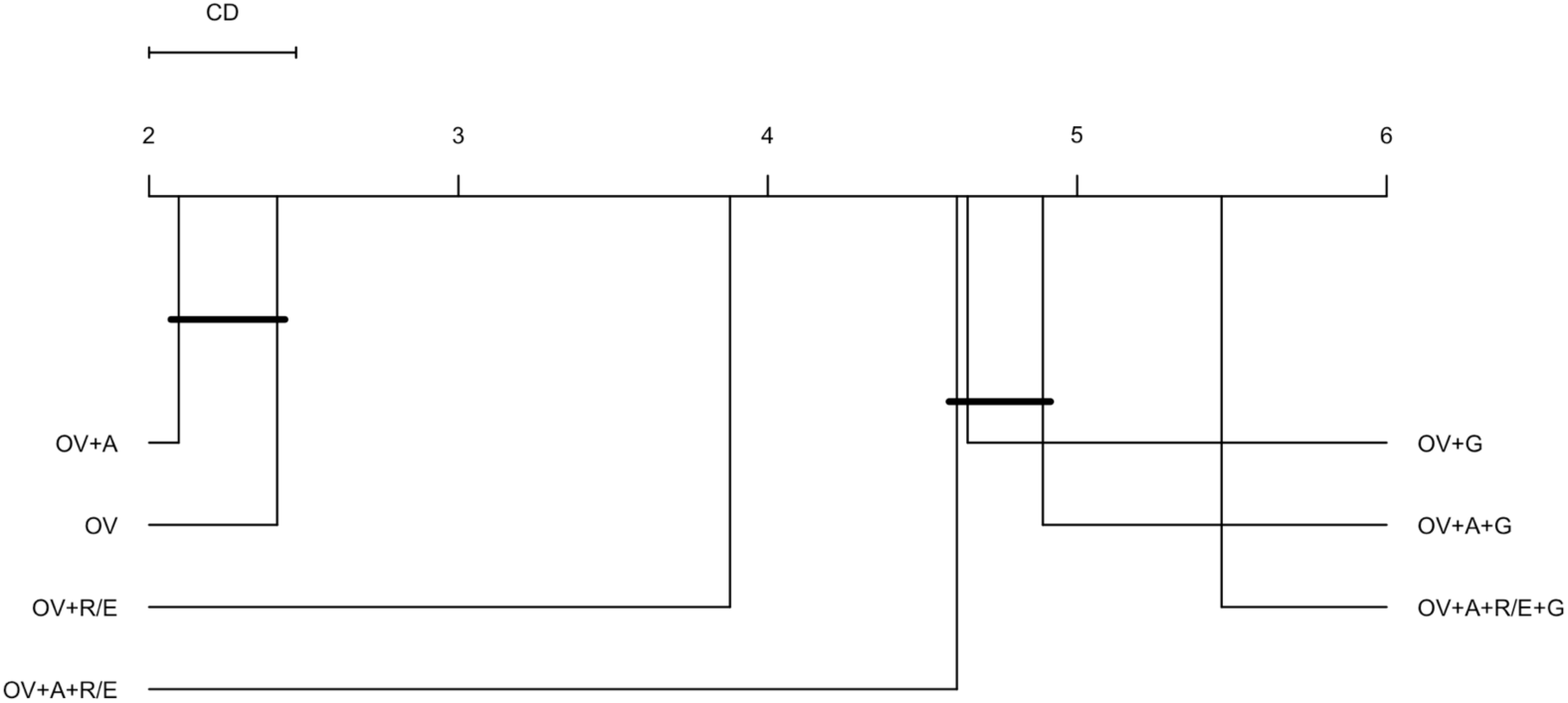
CD diagram of the average encoder performance rankings by features. * OV: observation value, A: age at observation, G: gender (male/female), and R/E: Race/Ethnicity (American Indian or Alaska Native, Asian, Black or African American, Native Hawaiian or other Pacific Islander, White Only, Multiple Race/Ethnicity, Hispanic, and unknown).

We obtained a critical difference of 0.475 on 7 different feature compositions and 2,513 degrees of freedom. We found that the encoders with two features, observation value (OV) and age at observation (A) scored the best overall outlier detection performance based on the Youden’d J index. The simplest encoders with only one feature – observation value (OV) – that was the runner up did not have a significant average performance ranking difference with the best encoders, at *p*-value < 0.05. Adding Race/Ethnicity (R/E) to the baseline encoder produced a better average performance ranking as compared with gender. We also found that the most complex combination of features – with all demographic variables included – performed the worst among all feature compositions. Friedman post-hoc test with Bergmann and Hommel’s correction also confirmed this ranking and significance testing.

So far, our results support our first hypothesis that adding demographic variables that already exist in most EHRs can improve the outlier detection performance – we only found that adding age at observation improves the average performance of our baseline encoders. To further evaluate the hypothesis, we evaluated the encoders by the combination of features and neural network (NN) architectures (Figure 4).

**Figure 4.**
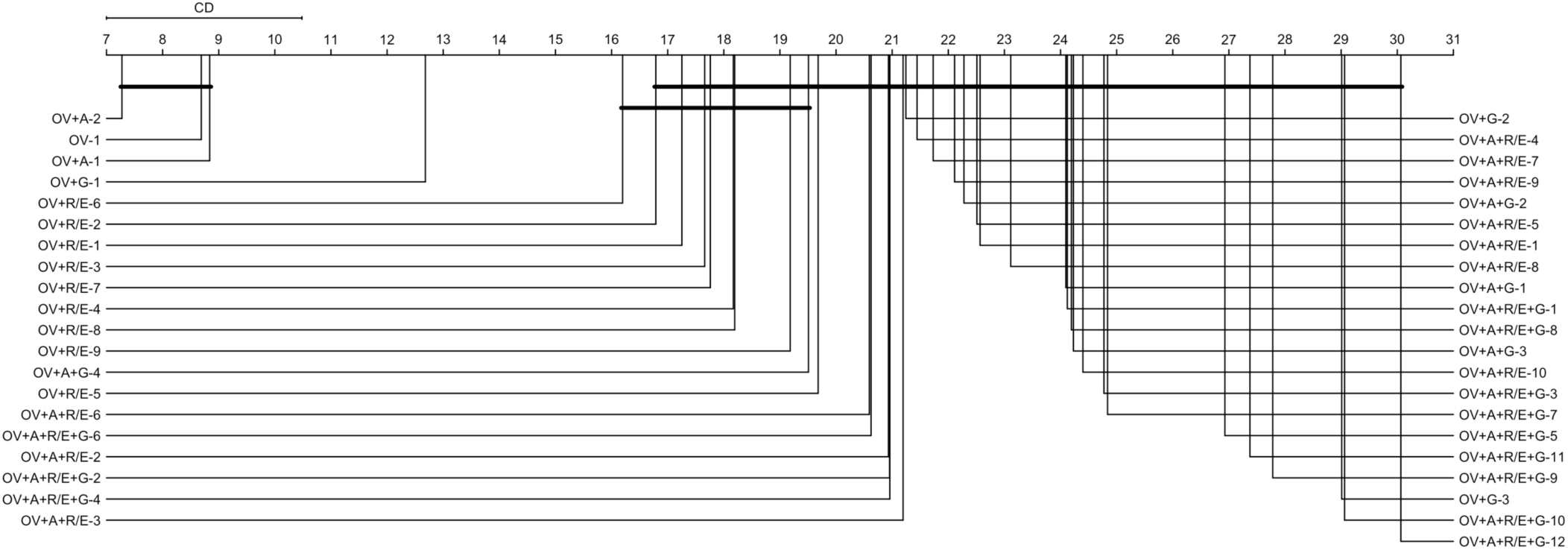
CD diagram of the average encoder performance rankings by the combination of features and NN architectures. * OV: observation value, A: age at observation, G: gender (male/female), and R/E: Race/Ethnicity (American Indian or Alaska Native, Asian, Black or African American, Native Hawaiian or other Pacific Islander, White Only, Multiple Race/Ethnicity, Hispanic, and unknown). The number after dash represent NN architecture – e.g., OV+A-2: encoder with OV+A as features and a 2⇒1⇒2 NN architecture.

Critical difference between encoders by the combination of features and NN architectures was 3.482, on 41 distinct feature-NN architecture combinations, and 14,719 degrees of freedom. Similar to what we found previously, still the encoders with two features, observation value (OV) and age at observation (A) scored the best overall outlier detection performance. We also found again that the most complex encoders, with all features and deepest NN architecture (12⇒11⇒10⇒9⇒8⇒7⇒6⇒5⇒4⇒3⇒2⇒1⇒2⇒3⇒4⇒5⇒6⇒7⇒8⇒9⇒10⇒11⇒12), produced the worst average outlier detection performance. In general, it appears that outlier detection performance decreased with adding complexity.

### Testing the second hypothesis

Each of the 41 distinct feature+NN architecture combinations (in Figure 4), we still have 240 combinations of sampling strategies and error margins. We first compared the average outlier detection performance ranking across the four different sampling strategies – 1%, 5%, 10% for semi-supervised encoders, and 100 percent in conventional autoencoders (Figure 5). With a critical difference of 0.077 and 14,756 degrees of freedom, results unanimously supported our hypothesis that using a sampling strategy would improve outlier detection performance. We found that semi-supervised encoders built on samples of 5 or 10 percent of data equally (at *p*-value < 0.05) produced the best average performances. The conventional autoencoders had the worst average outlier detection performance among the four sampling strategies. Friedman post-hoc test also confirmed this ranking and significance testing.

**Figure 5.**
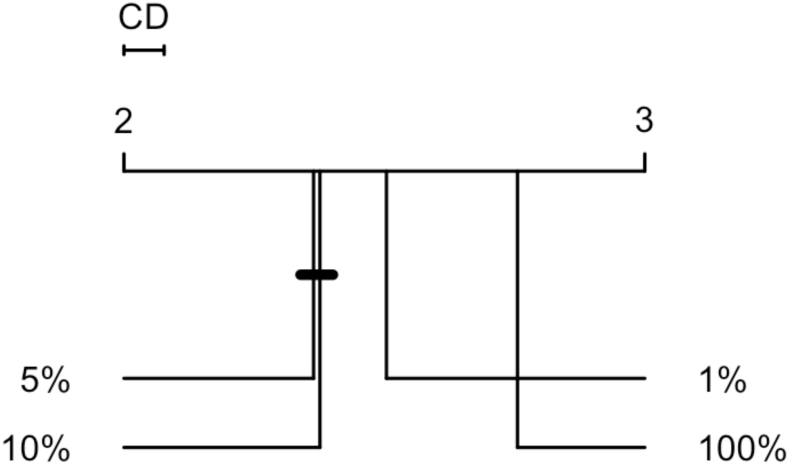
CD diagram of the average encoder performance rankings by the combination of features and NN architectures. * percentages represent sample-to-data ratio

## What is the best outlier detector?

After evaluating our hypotheses with non-parametric tests and calculating average performance rankings, it is taunting to see if an encoder consistently performed outstanding outlier detections across all 30 observations. Results showed that the best encoder performance for each observation has a minimum J index of 0.999 – i.e., there is at least an encoder for each observation that resulted in a J index better than 0.999. As the ROC plots showed (Figure 2), we expected performance variability between the 492 encoders and across the 30 observations. Figure 6 presents a box plot of J indices across all encoders. For the purpose of finding the best encoder, we focused on encoders that had a J index higher than 0.99 of the observations, and more importantly, also had low variability in high performance – such that their first quartile value was higher than 0.9. This left us with seven encoders (Figure 7 – also highlighted on Figure 6).

**Figure 6.**
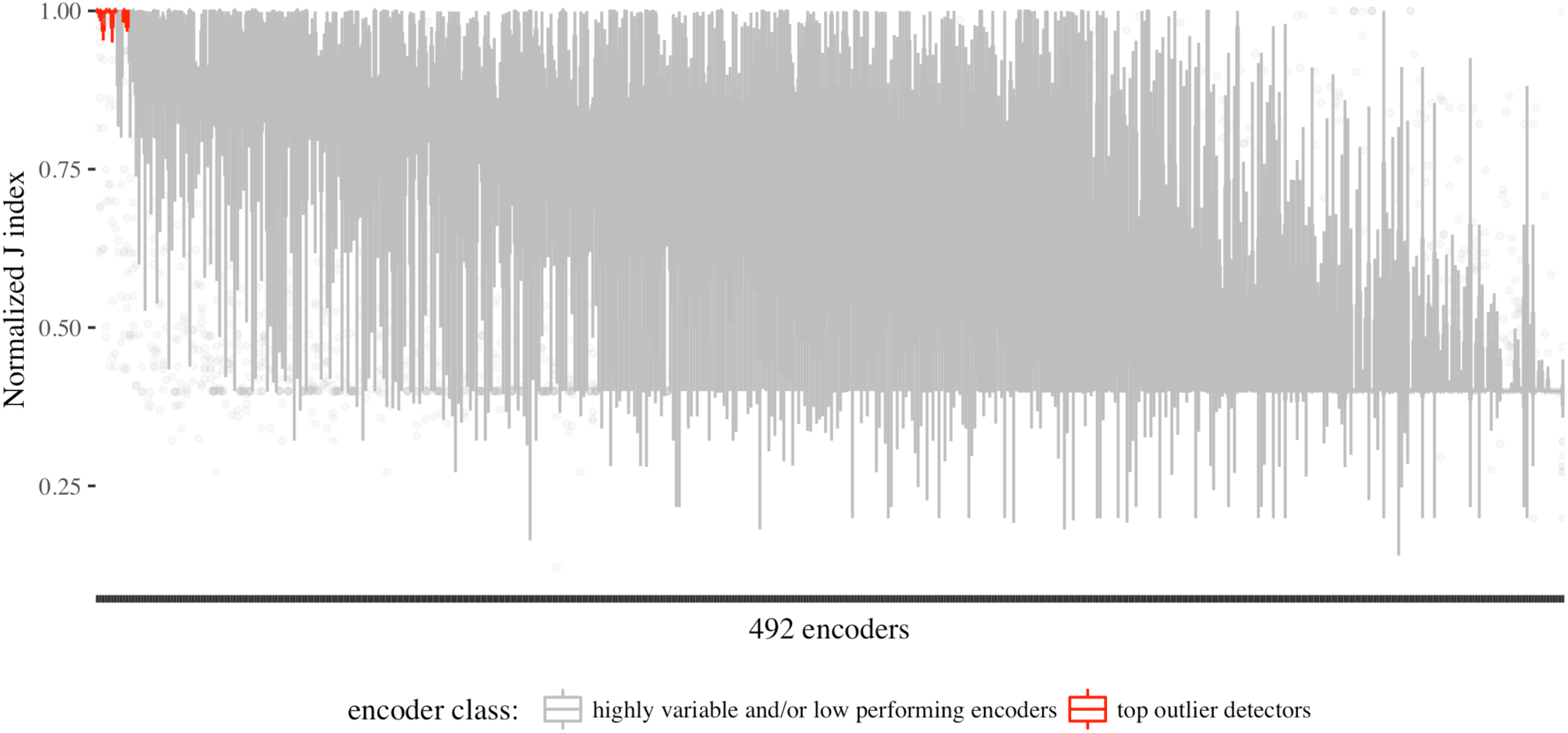
outlier detection performance variability across the 492 encoders.

**Figure 7.**
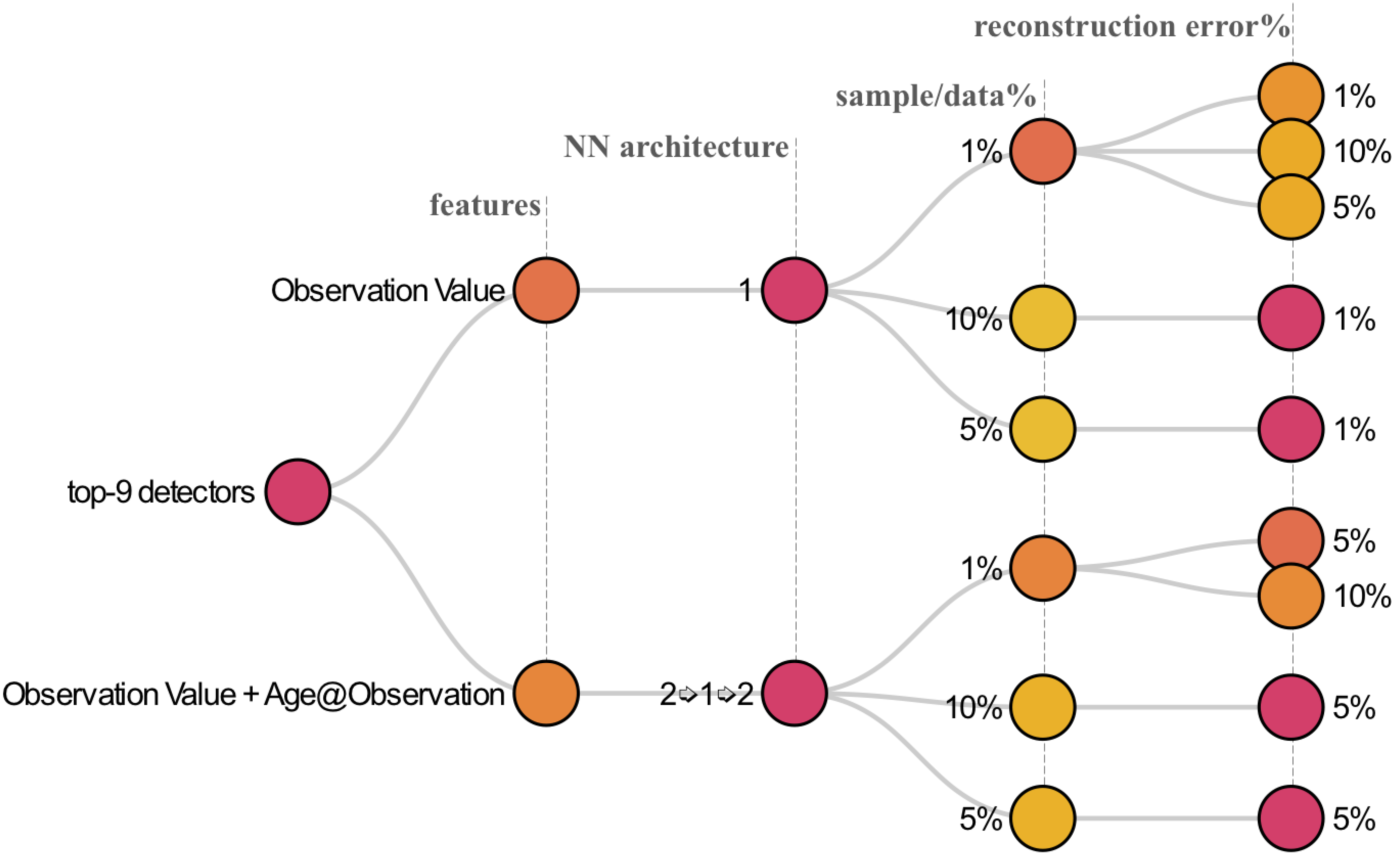
the top-9 outlier detector encoders based on the combination of performance and low variability.

The nine top outlier detector encoders comprised of 270 observations across the 30 observations, allowing us to also observe variability across observations. The box plots in Figure 8 show that while the top performing encoders frequently have reliably high performances across observations, there is only one encoder that performed well for BMI (LOINC 39156-5) and HCO3 (LOINC 1959-6) observations (shown as an outlier point), while most of the top encoders performed poorly for the two observations.

**Figure 8.**
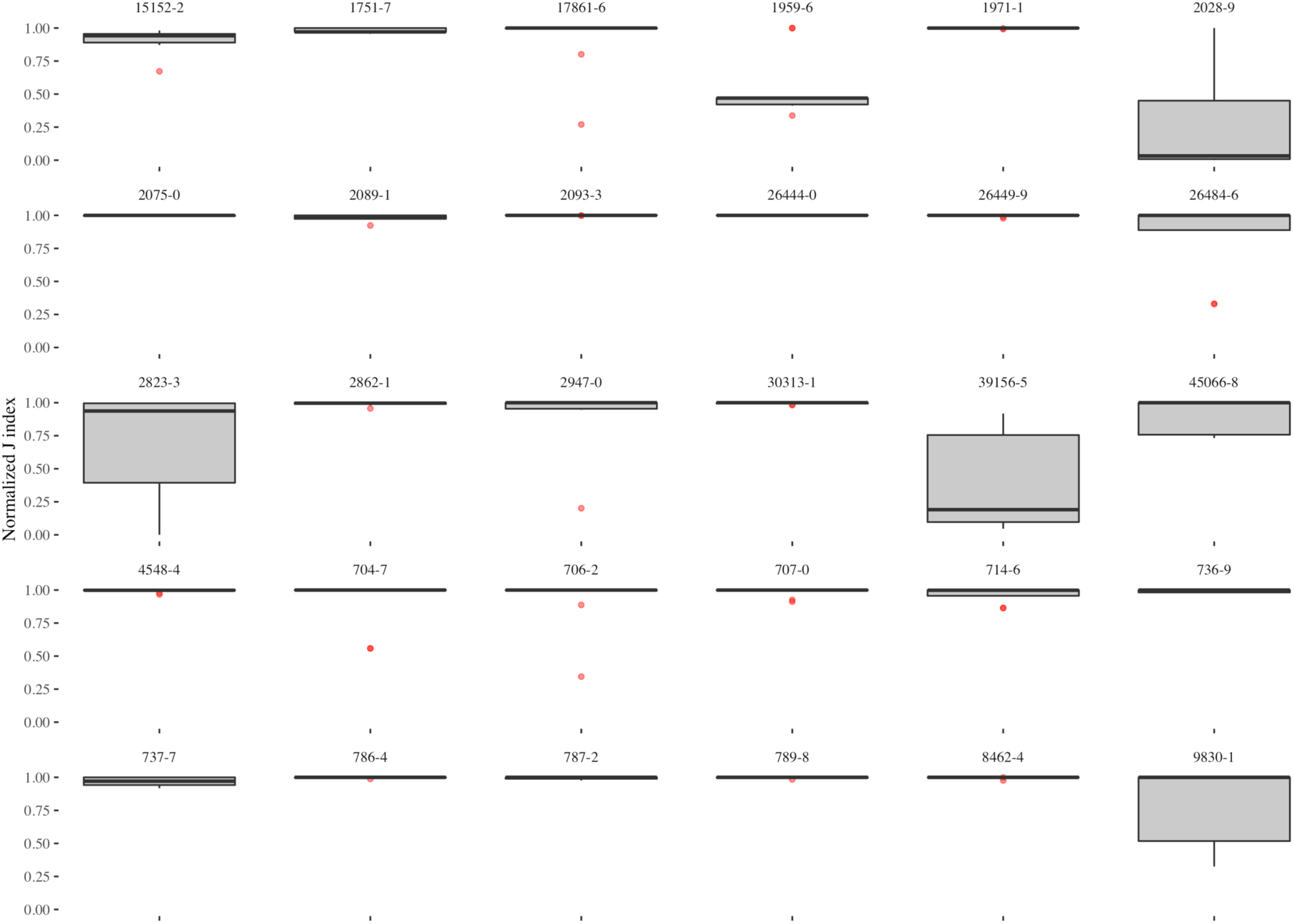
Variability of outlier detection performance in the top-nine encoders across the 30 observations.

To explore whether there is a common outlier detector encoder among the top-nine encoders that performed well across all observations we broke down J index for each of these encoders by observation. The heatmap on Figure 9 illustrates the normalized J index of the top-nine encoders across the 30 observations. As the figure shows, we found the best all-around encoder in a semi-supervised encoder with one feature – observation value (the baseline feature) – and a single hidden layer, which was built on one percent of the data and identified data points with more than one percent reconstruction error as outliers (OV-1-1%-1%).

**Figure 9.**
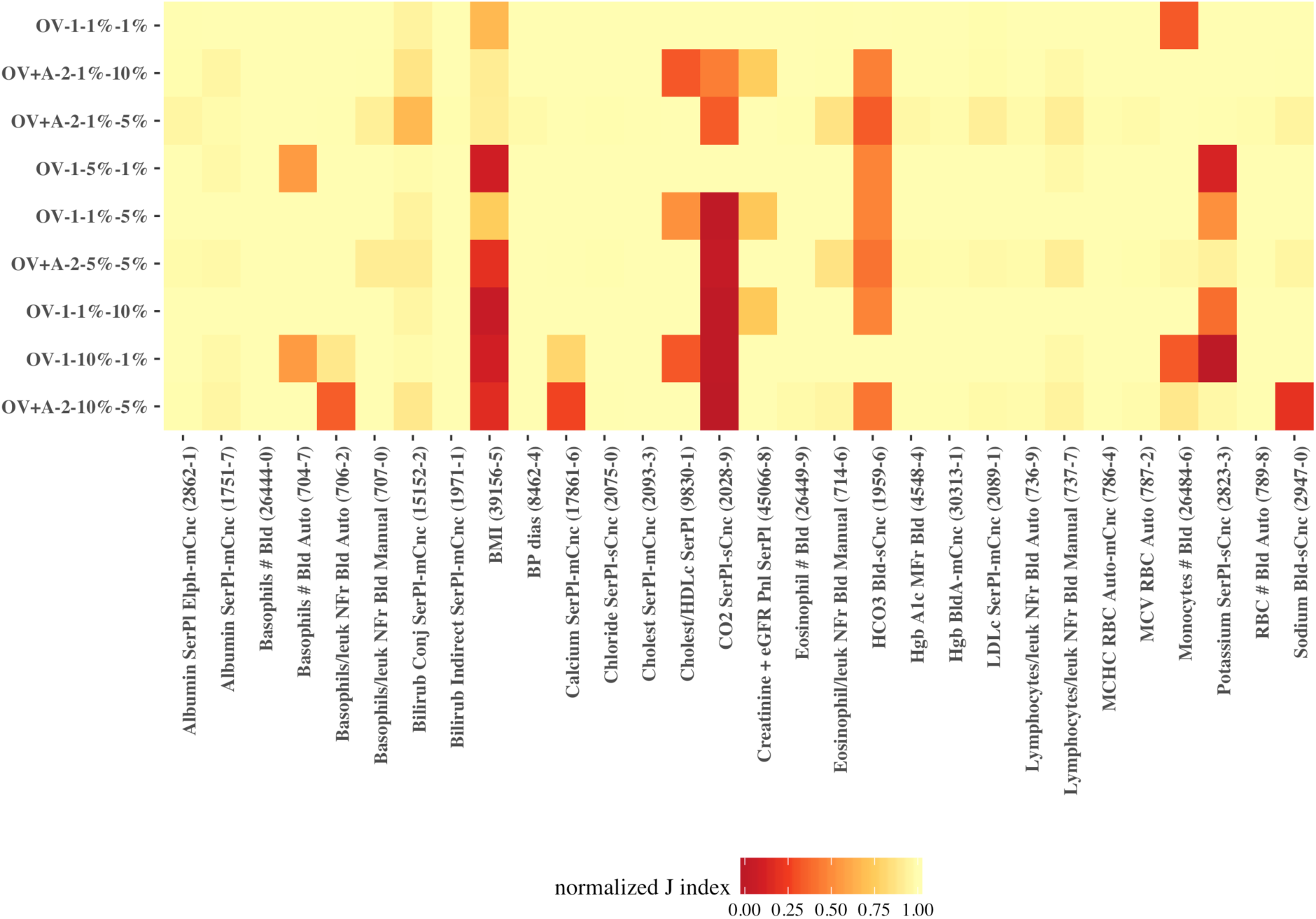
heatmap of the normalized J index for the top-nine encoders across the 30 observations.

To statistically verify these findings, we performed Friedman post-hoc test with Bergmann and Hommel correction on the top-nine encoders (Table 1). Applying Bergmann Hommel correction to the *p*-values computed in pairwise comparisons of nine algorithms requires checking 54,466 sets of hypotheses. The Friedman post-hoc test shows that the top-performing encoder was not significantly different (at *p*-value < 0.05) from semi-supervised encoders with similar NN architecture (single hidden layer) and feature (observation value), which applied different sampling strategies and error margins (5% or 10%). The top-performing encoder was also statistically not different from the encoder that used the same sampling strategy (one percent) but had age at observation and observation value as features (OV+A) in a 3-layer (2⇒1⇒2) NN architecture. Nevertheless, the top encoder’s performance was higher for all 30 observations than the two that generally were not significantly different.

**Table 1.** Corrected Friedman post-hoc *p*-values * insignificant p-values (< 0.05) with Bergmann and Hommel correction are highlighted in Bold for pairwise similarities. The top-performing encoder is highlighted first row of the matrix.

| *feature* | *OV* | *OV* | *OV* | *OV* | *OV+A* | *OV+A* | *OV+A* | *OV+A* |
| --- | --- | --- | --- | --- | --- | --- | --- | --- |
| *architecture* | *1* | *1* | *1* | *1* | *2* | *2* | *2* | *2* |
| *sample ratio* | *1%* | *1%* | *10%* | *5%* | *1%* | *1%* | *10%* | *5%* |
| *error margin* | *10%* | *5%* | *1%* | *1%* | *10%* | *5%* | *5%* | *5%* |
| *OV-1-1%-1%* | **0.33** | **1.00** | **1.00** | **1.00** | **0.76** | 0.00 | 0.00 | 0.00 |
| *OV-1-1%-10%* |  | **1.00** | **0.63** | **1.00** | 0.00 | 0.00 | 0.00 | 0.00 |
| *OV-1-1%-5%* |  |  | **1.00** | **1.00** | **0.14** | 0.00 | 0.00 | 0.00 |
| *OV-1-10%-1%* |  |  |  | **1.00** | **0.43** | 0.00 | 0.00 | 0.00 |
| *OV-1-5%-1%* |  |  |  |  | **0.10** | 0.00 | 0.00 | 0.00 |
| *OV+A-2-1%-10%* |  |  |  |  |  | **0.05** | **0.33** | **0.33** |
| *OV+A-2-1%-5%* |  |  |  |  |  |  | **1.00** | **1.00** |
| *OV+A-2-10%-5%* |  |  |  |  |  |  |  | **1.00** |

The top-nine encoders on average had a J index of 0.9968 for the 30 observations, with a standard deviation of 0.0143. The minimum best performance of the top nine encoders across all observations was 0.9230 – for BMI (LOINC: 39156-5). However, at least one encoder, which may be outside the top-nine encoders had a Youden’s J index higher than 0.9999 for all 30 observations. For example, the top-two performances for BMI had J indices of 0.99998 and 0.99994, respectively from OV+G-1-1%-5% and OV+A-1-1%-5% that were not included to the general top models.

## Discussion

Predictive (a.k.a., unsupervised) learning techniques present novel low-cost opportunities for detecting complex patterns in unlabeled clinical data. Given the availability of computing power and large scale clinical datasets, encoders are effective methods for compressing non-linear representations of clinical data. We demonstrated the utility of semi-supervised encoders (super-encoders) for outlier detection through compressing the data into an exemplar distribution learned from hypothetically less-noisy random samples. In addition to precision, encoders are often very quick in compressing the data. On average, it took 7.034 seconds for the top seven encoders to compress a dataset with an average size of over 7 million rows. Taking into account the multiplicity of distributions for a single observation in EHR data (i.e., same observation represented with different names and/or units), encoding offers huge promises for outlier detection in large scale clinical data repositories.

Generally, we found that simple super-encoders produced outstanding outlier detection performances. We were able to find a super-encoder that performed very well for all 30 observations. The top encoder only used the observation value and a single hidden layer with one neuron to compress and decompress the data – essentially a simple regression model with tanh as the activation function.

We also found that only adding age at observation to the baseline encoder (observation value) improved outlier detection. However, we used a linear cut-off range (based on observation value) to measure the performance of each encoder. Defining ranges in multiple dimensions (which would form a hyperplane) is needed to further evaluate the performance of encoders. Nevertheless, even the linear range definition proved encoding as a feasible alternative for replacing the current manual standard procedures for outliers (i.e., identifying biologically implausible values) in clinical observations data. We also found that the more complex an encoder gets (i.e., including more demographic features and increasing depth of the NN), the smaller its reconstruction error is. This is in agreement with prior research that holding the number of features constant, a deep autoencoder can produce lower reconstruction error than a shallow architecture.[13] In other words, the compressed data representation produced by a deep NN encoder with all demographic features was very close to the actual data, which included outliers. This finding provides further evidence for the potential utility of denoising encoders in phenotyping and compression of clinical data.

Looking forward, with the availability of state-of-the-art computational resources and increasing amounts of clinical data in today’s healthcare organizations, training generalizable super-encoders for each group (or combination of groups) of clinical observation data seems feasible. Our semi-supervised approach takes the view of letting the data speak for itself and can improve (or at least provide an alternative to) the current rule-based approaches to identification of implausible values that often do not take non-linearities into account.

The experimental work conducted in this research was on 30 observations. Therefore, our comparison and ranking of the algorithms – which resulted in identifying the top encoders – is based on validation results from a limited number of observations. For improving generalizability of these findings further efforts may be needed to include validation results for a larger set of observations. We envision the top encoder (or a combination of the top performing encoders) to operate on the data warehouse after each data refresh. Once the outliers are detected and flagged, a workflow that would likely involve context expertise is needed to validate the flagged observations and initiate further actions.

## Conclusion

We found a best all-around encoder; a semi-supervised encoder, with observation value as the single feature and a single hidden layer, trained on one percent of the data, identifying outliers as data points with higher than one percent reconstruction error. The top-nine encoders on average had a J index of 0.9968 for the 30 observations. At least one encoder, which may be outside the top-nine encoders had a Youden’s J index higher than 0.9999 for all 30 observations. Due to multiplicity of observation data and their representations in EHRs, detecting outliers (i.e., biologically implausible observations) is challenging. Even in the presence of gold/silver standard implausibility thresholds, non-linearities of human observations are not captured in linear cut-off ranges. We demonstrated the utility of a semi-supervised encoding approach (super-encoding) for outlier detection in clinical observations data. Super-encoder functions and their implementations on i2b2 star schema is available on GitHub (*link will be posted after the peer review process*). In addition, analytic scripts to perform the experiments and analyze the results in R statistical language are available on GitHub for reproduction of the experiments across other EHR data warehouses.

